# Interfacial energy constraints are sufficient to align cells over large distances

**DOI:** 10.1101/653535

**Authors:** S. Tlili, M. Shagirov, S. Zhang, T. E. Saunders

## Abstract

During development and wound healing, cells need to form long-ranged ordered structures to ensure precise formation of organs and repair damage. This requires cells to locate specific partner cells to which to adhere. How such cell matching reliably happens is an open problem, particularly in the presence of biological variability. Here, we use an equilibrium energy model to simulate how cell matching can occur with subcellular precision. A single parameter – encapsulating the competition between selective cell adhesion and cell compressibility – can reproduce experimental observations of cell alignment in the *Drosophila* embryonic heart. This demonstrates that adhesive differences between cells (in the case of the heart, mediated by filopodia interactions) are sufficient to drive cell matching without requiring cell rearrangements. The biophysical model can explain observed matching defects in mutant conditions and when there is significant biological variability. We also demonstrate that a dynamic vertex model gives results consistent with the equilibrium energy model. Overall, this work shows that equilibrium energy considerations are consistent with observed cell matching in cardioblasts, and has potential application to other systems, such as neuron connections and wound repair.

**Statement of Significance:** Cells often need to identify specific neighboring cells, such as during wound repair and forming neural connections. Here, we develop a biophysical model of such cell-cell interactions within the context of the developing heart. We demonstrate that precise cell matching can occur by minimizing the energy costs of interfacial interactions. This model can explain a breadth of experimental observations despite it being a steady-state approximation of a dynamic system. This opens the possibility that such approaches may be applicable to other systems, providing a powerful yet simple framework for understanding cell matching.

## Introduction

During development, cells interact collectively to form tissues and organs through a series of morphological transformations, driven by cell proliferation, rearrangements, migration and death (1-5). When these processes fail, the final shape of the tissue can be defective, resulting in diseases, including cardiomyopathies (6, 7) and neurological defects (8). During organogenesis, cells often need to identify specific cells to which to adhere (9). A classic example is formation of human facial structures (8, 10); cells initially undergo long-ranged migration from distinct regions of the neural plate before forming precise connections to create structures such as the lip. Errors in this process lead to birth defects such as cleft lip and facial cleft (10). During neurogenesis, neurons also need to form precise linkages to their synaptic partners (11-13) with severe consequences if these processes fail. A range of molecules have been identified that are involved in cell matching, predominantly from neuronal systems (14-17). These molecules include cytoskeletal, adhesion, and force transducing proteins (9). However, the underlying mechanisms by which the information from these different components is integrated by the developing tissue to form precise connections remain unknown.

In most tissues, there are multiple cell types with stereotypic spatial positions. For example, formation of the eye requires precise cell fate determination and positioning (18, 19), even within a growing domain (20). In the developing heart, cardioblasts take on different fates depending on expression of (highly conserved) transcription factors (21). Periodic patterns of cells can also be generated from lateral inhibition (22, 23). Such periodic patterns need to be maintained across large distances, even as tissues undergo large-scale morphological changes. Experimental and theoretical work has begun to reveal how neighboring tissues can form distinct boundaries (24, 25). An important concept in generating and maintaining tissue structural organization is the Differential Adhesion Hypothesis (26, 27). By cells mechanically interacting differently with each other, depending on specific fate, cells can form ordered structures and patterns.

Theoretical modelling of tissue formation has helped increase our understanding of how cells pack (28), form compartment boundaries (29), generate complex tissue shapes (30-34), and ensure regulated growth (35). Lateral inhibition can create a wide variety of patterns depending on the feedback mechanisms (36). Recently, vertex models have been used to understand cell structure in epithelia (37, 38), including in three-dimensions (39). Although biological systems are inherently dynamic, equilibrium statistical mechanics can be a powerful tool for understanding suitable biological processes. An example of such a case is cell packing in the eye, where analogies with soap bubbles provide an effective tool for understanding formation of the *Drosophila* retina (40).

Here, we develop a model of cell matching, where differential adhesion energy constraints between cells drives the process of matching. We apply this to the developing *Drosophila* embryonic heart, where the process of cardioblast cell matching has been quantified (41-43). The *Drosophila* embryonic heart is comprised of two lines of cardioblasts that migrate together over a period of a few hours, Fig. 1A, and they express either Tinman (Tin, the *Drosophila* homolog of mammalian Nkx2.5) or Seven-up (Svp), Fig. 1B, in a repeating 4-2 pattern (44). Cardioblasts first approach each other through migration towards the embryo midline, driven by surrounding tissues (45) and myosin contractility (46), which we refer to as a *ballistic* phase, Fig. 1C. As cardioblasts approach each other, the cells adjust position, via filopodia interactions between cardioblasts, to align accurately with their contralateral partners, Fig. 1D. Specific adhesion molecules are expressed within the different cardioblast types: Fasciclin III (Fas3) in Tin-positive cardioblasts; and Ten-m in Svp-positive cardioblasts, Fig. 1E. Cell matching in the heart depends on the differential spatial expression of these adhesion molecules (41). The mechanical force between opposing cells is generated by a periodic wave of Myosin II in the cardioblasts, that acts to “proof read” the mechanical connections (42, 43). Motivated by these observations, we construct a biophysical model of the cell-cell interactions to test whether such differential adhesion is sufficient to drive cell matching. We then use this model to test how both structural and genetic perturbations alter cell matching. This model provides a biophysical framework in which to understand how cells within confluent tissues find specific partners during development.

**Fig. 1.**
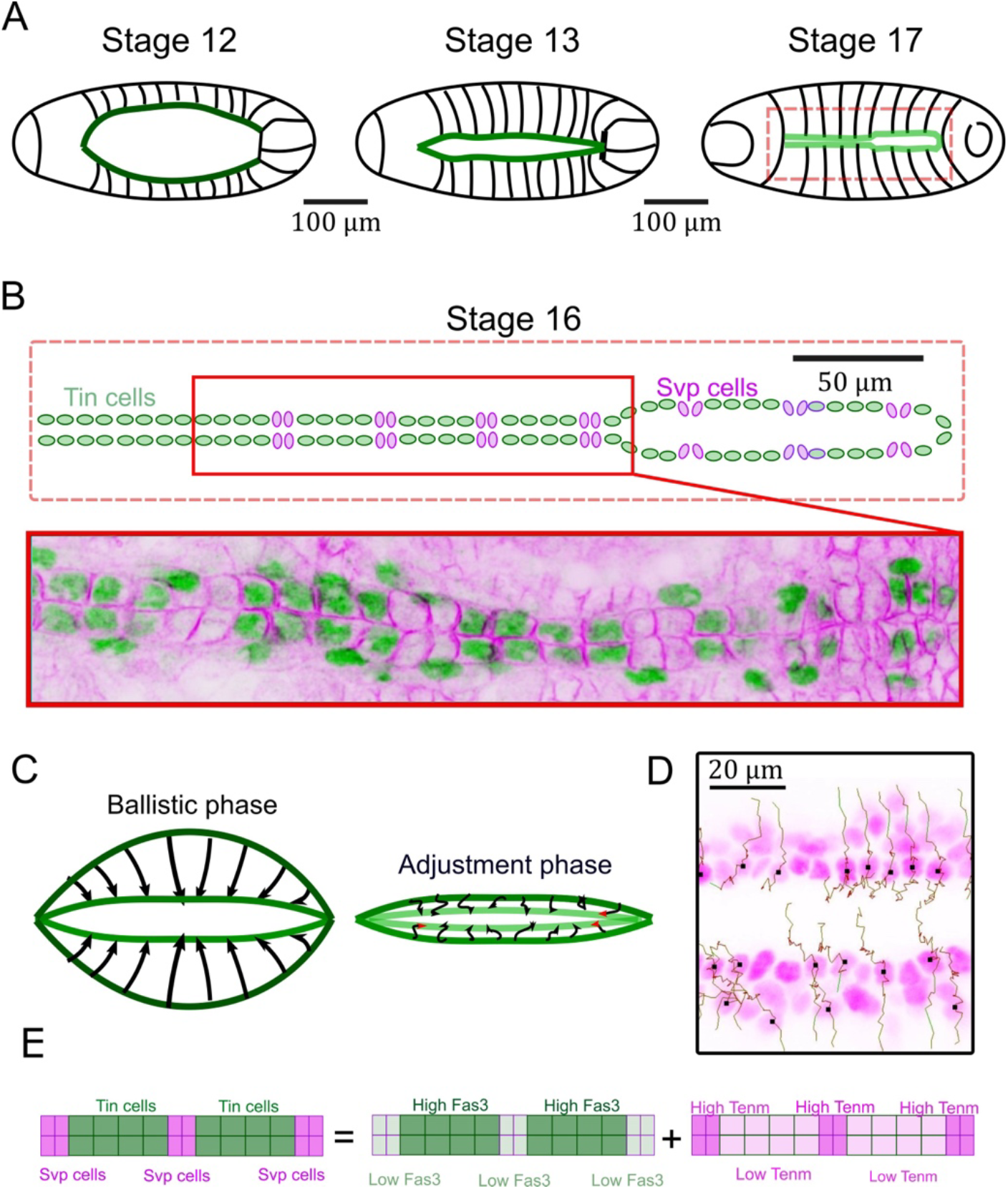
Cell matching in *Drosophila* embryonic heart. (A) Developmental stages of heart formation. Cardioblasts (green regions) merge to form a tubular heart (stage 17). (B) Cardioblasts have distinct expression patterns. Heart structure at stage 16: the heart is made of a periodic alternation of Tin cells and Svp cells. Staining of cardioblasts (Spectrin in magenta and Tinman in green). (C) Heart formation is initially driven by a global tissue movement (a ballistic phase in which cardiac cells are passively driven by the dorsal closure process). Once the two contralateral rows of cardioblasts are almost in contact, active processes align cells in a final adjustment phase. (D) Example of cell trajectories during the ballistic (straight trajectories) and adjustment phase (diffusive trajectories). (E) Cardioblasts express the adhesion molecules Fas3 and Ten-m in an alternating pattern.

## Materials and Methods

### Methods

Experimental data shown is taken from (41, 42). A detailed methodology for the quantification of heart cell matching is available (43).

### Modelling

#### One-dimensional interface, and mismatch ratio definitions

The model system consists of two rows of equal number of cells interacting on a one-dimensional interface defined as vertices *x*, Fig. 2A. Each cell *i* on a row *r* is represented by a pair of vertices 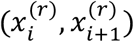 on this interface with 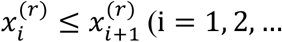, N +1, and r ∈{1, 2} for N cells occupying each row), with cell length

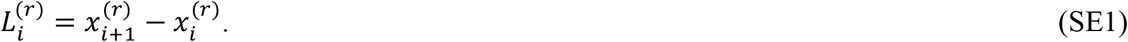

**Fig. 2:**
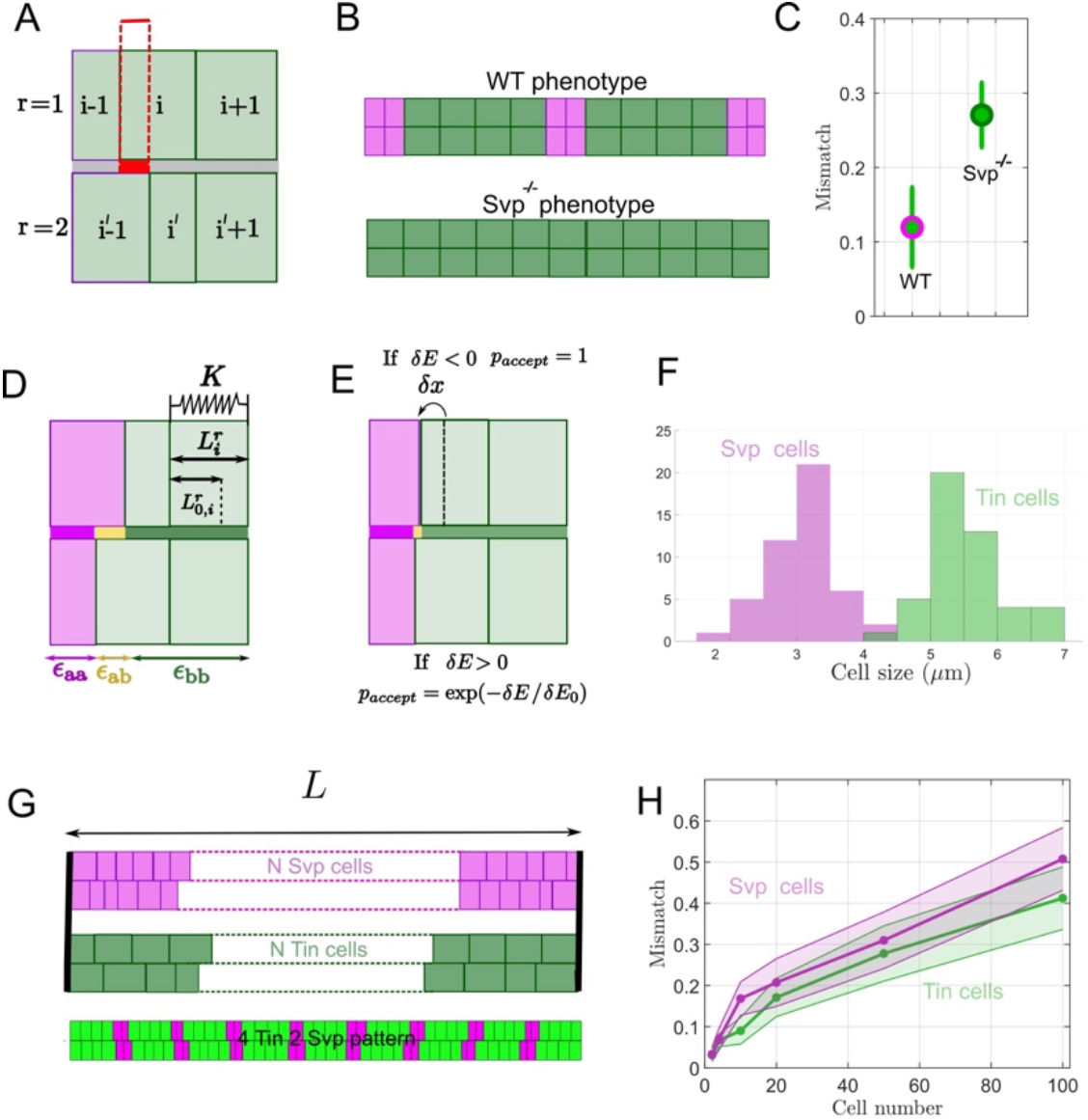
Simulating heart matching. (A) Definition of cell mismatch. *r=1, 2* denotes two contralateral rows of cardioblasts. The dashed red box denotes a “mismatched” region in which two cells which are not supposed to be in contact are partially facing. (B) The wildtype heart (top) has a stereotypic pattern of cardioblast fates (denoted by magenta (Svp positive) and green (Tin positive) cells), which express specific adhesion molecules. Such a pattern is lost in *svp* ^*-/-*^ mutants (bottom), where there is a single cardioblast type, Tin cells. (C) It has been experimentally shown by some of us that heart cell mismatch (as defined in Fig. 2A) is affected by the loss of adhesion pattern in *svp* ^*-/-*^ mutants (41). (D) Considering two cell types (denoted by *a* (magenta) and *b* (green)) there are three interface energies: ε_aa_ between two cells of type a, ε_bb_ between two cells of type b, and ε_ab_ between cells of different type (denoted by yellow region). Cells can deform elastically (or compress), *L* _*i*_ from their rest length, *L* _*0*_ with a compressibility K and cell-cell contact is characterized by an adhesion depending on the contact type. (E) To find how cell differential adhesion competes with the elastic deformations generated by local cell compression, a metropolis algorithm is used to find the final cell alignment after equilibration. At each step, a cell/cell interface is randomly chosen and is displaced by δx, which follows a Gaussian distribution (see Methods). The probability to accept the displacement follows the Metropolis algorithm. (F) Experimental distribution of leading-edge size in different cardioblast types (41). (G) Simulations with only one cell type (all Svp or all Tin) and with the alternating pattern of Tin/Svp cells (bottom). (H) Predicted mismatch in hearts with one cell type for different cell number. The mismatch increases with the linear size of the system (the number of cells in a heart row) as each cell width varies as a random variable.

The total length of a model system is kept constant by fixing first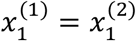, and last 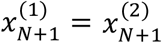vertices throughout the simulation. The overlap interval length between cells *i* and *j* located on two different rows is calculated as:

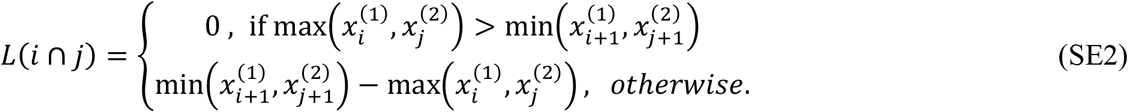

The mismatch ratio 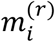 of a cell *i* on a row *r* is defined as a proportion of length of overlap with cells other than its sister cell *i* ′, and the total cell length of *i* :

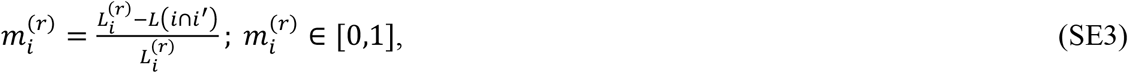

where the sister cell is defined as a cell *i* ′ with same index as *i* but located on a different row (i.e. *i* ′ = *i*), an average mismatch ratio of two sister cells is then calculated as 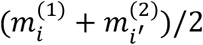.

#### Calculating net change in energy

All vertices 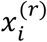 are assumed to be coupled with harmonic springs, with each cell having an elastic energy 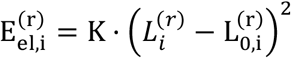,where 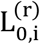 is the equilibrium length for a given cell (kept constant throughout the simulation), and K is a spring constant. Then, the total elastic energy for the whole system is just a sum of elastic energies over all cells:

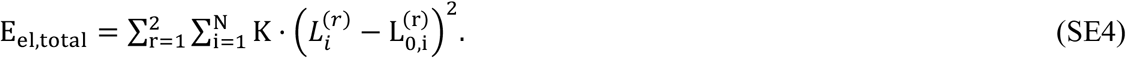

In the model, two rows of cells interact along the one-dimensional interface by adhesion of cells along segments of the interface shared by the same cell type cells, or by cohesion of cells of two different cell types along interface segments occupied by cells of two different types.

We implemented the interface by defining interface vertices 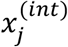 with 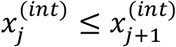, which could be obtained by concatenating both rows of 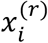 vertices into single vector and then sorting them. Similar to cell vertices 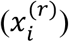, pair of interface vertices 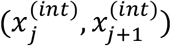 represents an interface segment *j*. The total adhesion energy for the whole system is then the sum of adhesion energy contributions of all interface segments:

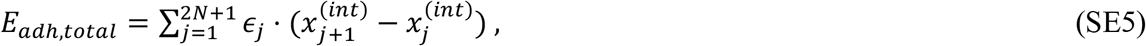

where ϵ_*j*_ is the cell type specific adhesion energy-per-unit length of a segment *j* :

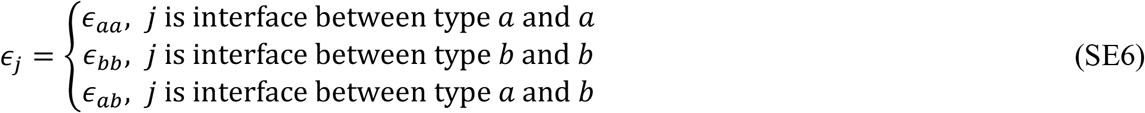

Since vertices of 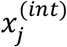 are from both rows of cells, changing position of a single cell vertex 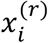 could result in cases with 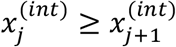, thus 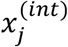 needs to be sorted before calculating total adhesion energy during simulation.

The total energy of the system as a function of location of cell vertices 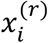 is calculated as the sum of elastic and adhesive energies:

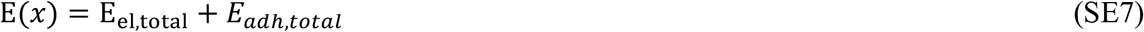

and the net change in total energy due to change in the configuration of *x* → *x* +Δ*x* is calculated as a difference in total energy between the two configurations

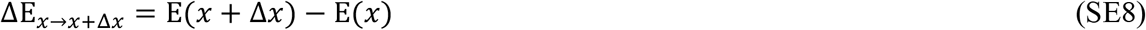

#### Sampling initial configuration and equilibrium lengths of cells

Randomness in cell shape geometry was modelled as a random initial configuration of cells, which was set to be equal to the equilibrium lengths of the cells throughout the simulation. The random initial configuration of cells, and thus the equilibrium lengths 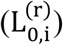 for each cell were sampled from a normal distribution with cell type specific mean and standard deviation using a MATLAB function *normrnd*. In order to construct a random configuration, we implemented random sequential deposition (47) of two rows onto each other with a constraint that enforces equal length of two rows within ε = μ ⋅ 10^−3^, where μ is the smallest of the two type-specific average lengths of the cells in the given configuration.

For the row of cells with m_a_ type a cells and N – m_a_ type b cells, m_a_ equilibrium lengths 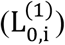 for the top row are sampled from a normal distribution with mean 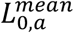 and standard deviation 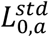, and N – m_a_ equilibrium lengths 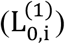 are sampled from normal distribution with mean 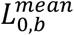 and standard deviation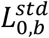.

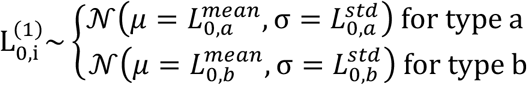

where the mean and standard deviation for different cell types are experimentally determined parameters. Next, the bottom row equilibrium lengths 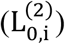 were sampled in the same manner until the constraint 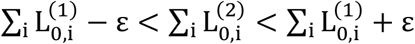 was satisfied. Afterwards, 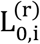 are converted into cell vertices 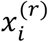 in the same sequence as the 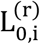 was produced by *normrnd* function.

#### Implementation of Markov-chain cell configuration sampling using Metropolis algorithm

In order to simulate evolution of the cell boundaries by moving cell vertices *x* and sample cell configurations, we used Monte Carlo Markov-chain (MCMC) sampling by implementing Metropolis algorithm in MATLAB (48). At each MCMC sampling step a vertex 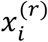 is selected randomly and a random move is proposed 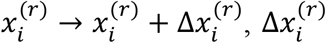 is a Gaussian random variable with mean 0 and standard deviation δ*X*, the value of δ*X* determines magnitude of the random movement (interpreted as random fluctuations in the model). In our implementation 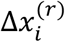 was sampled using *normrnd(0*, δ*X)* function in MATLAB. Afterwards, Metropolis probability of accepting the proposed move, 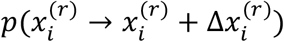 is calculated using

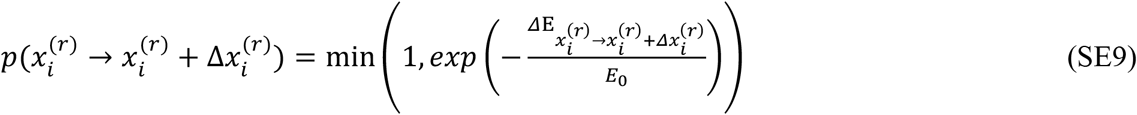

where 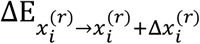 is a net change in total energy of the system due to movement of vertex 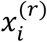 by amount 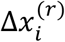, and *E* _0_ is the effective temperature (“thermal energy”) of the cells. Then, 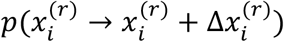 is compared to a uniformly distributed random variable δ_*p*_ in the interval δ_*p*_ ∈ (0,1) produced by *rand* function in MATLAB. If the statement 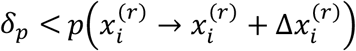 is true, then the move is accepted, and otherwise the move is rejected.

Overall, the probability of movement 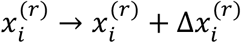 of a vertex at position 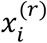is:

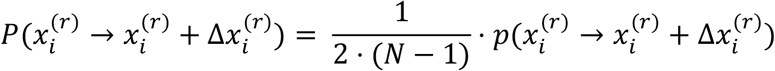

where the first term on the right-hand-side refers to the probability of selecting vertex 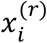 from two rows, and *N* − 1 vertices in each row (two end vertices on each row are assumed to be fixed throughout the simulation, and have *P* = 0). In order to avoid flipping of cells, and overlapping of cells on a single row, we forbid such movements by setting 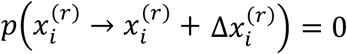 for movements with 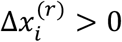 that exceed cell length of cell *i* (i.e. 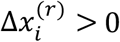 AND 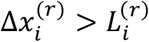, and for movements with 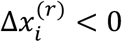 that exceed cell length of cell *i* – 1 (i.e. 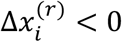 AND 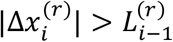).

#### Implementation of vertex model

Cell initial geometries are constructed using the same functions as with the equilibrium energy model above. Each vertex is described by its position 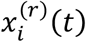,and follows the equation:

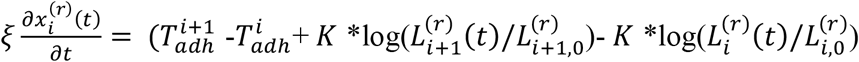 with *T* _*adh*_ = *T* _*SS*_ for a S-S interface, *T* _*adh*_ = *T* _*TT*_ for a T-T interface, and *T* _*adh*_ = *T* _*ST*_ for a S-T interface. Here, we use the elastic force form 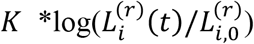 instead of 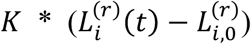 to avoid numerical artefacts at big cell deformation. These two definitions are the same for small cell deformations. Time is discretized in time steps dt = 0.01 min and the parameter ξ order of magnitude is tuned to observe the dynamics of the system evolution on the minute/hour timescale. Our previous equilibrium model predicts that matching will occur when *K* ∽ *T* _*adh*_ and on the hour timescale when *T* _*adh*_ /ξ ∽ 1 μ*m* / *min*. In Fig. 6, *T* _*adh*_ and *K* are expressed in pN and ξ in pN *min / μ*m*. The vertices positions are calculated at each simulation step and allow to calculate the mismatch evolution in time.

## Results

### Mapping of adhesive interactions between filopodia onto an equilibrium energy state

To simulate cell matching, we utilized our energy-based model (Methods) that accounts for the spatial constraints between cells and the adhesion competition between different cell types. In the *Drosophila* heart, differential adhesion between cells is mediated by filopodia contacts, which have a spread of contact times (41, 42), from a few seconds to over five minutes. As justified below, the underlying principle of the model is that these differential filopodia adhesion times between adhering cells can be mapped onto effective adhesion energies.

At a biochemical level, the time for reactions, τ, are typically related to the Arhennius law: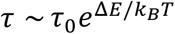, where Δ*E* is the energy cost of the reaction. Time scales in biological systems at macroscopic levels often follow the Arhennius law, such as in the developmental time of *Drosophila* embryogenesis (49, 50). Here, we apply a *mesoscopic* approximation, where we assume that the distribution of binding times of filopodia τ_*bind*_ can be related to the effective adhesion energy barrier Δ*E* _*adh*_ to separate contacting filopodia.

In the model, the overlap between facing cells results in an adhesion energy Δ*E* = −ϵ. *x*, where *x* denotes the length of the cell contact interface and ϵ is the adhesion energy per unit length between the two cells. In previous work (41), we showed that the expression of specific adhesion molecules is patterned along the heart, Fig. 2B; disruption to this pattern leads to matching defects, Fig. 2C. We can encode these adhesion differences as effective energy in our model, Fig. 2D. Cellular compressibility for each cell is encoded by an effective elastic energy *E* _*el*_ = *K* (*L* _*cell*_ – *L* _0_)^2^, corresponding to the cost of deforming the cells away from their preferred cross-sectional width; *K* is the effective cell compressibility along its leading edge, *L* _*cell*_ the cell length and *L* _0_ the cell rest length, Fig. 2D. We only focus on the apical leading edge of the cells as this is where the process of cell alignment is occurring. Combining the energy scales, we can express effective energies for two cell types, denoted by *a* and *b* :

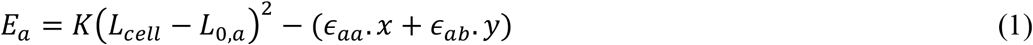

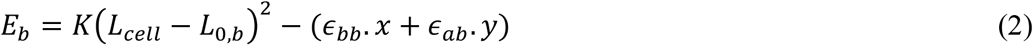

ϵ_*ab*_ denotes the adhesion energy per unit length between cells of type *a* and *b, x* and *y* denote the total alignment overlap with cells of the same and different types respectively, and *L* _0,*a*_ and *L* _0,*b*_ represent the equilibrium lengths for the different cell types.

Filopodia activity results from the interplay between active fluctuations and adhesion interactions with other filopodia. The active alignment of the heart takes place over a period of around 30 minutes, whereas the average binding time of filopodia is 1-5 minutes. In the following, we assume that the heart has enough time during the active alignment process to reach equilibrium.

To simulate the evolution and equilibration of configurations of heart cell, we use a Metropolis algorithm incorporating mechanical fluctuations induced by filopodia activity as an effective temperature, see Fig. 2E and Methods for further details. We define cellular mismatch by identifying the fraction of cell boundaries that are not correctly aligned with their corresponding opposite cell, Fig. 2A. In this definition, a perfectly aligned tissue has mismatch of 0, while a fully misaligned system has a mismatch of 1. The tissue mismatch is then taken as the average over all cell mismatches.

### Minimization of interfacial energy constraints are sufficient to explain cell alignment in the wildtype *Drosophila* heart

We apply our model to the developing *Drosophila* heart, where quantitative data for cell matching is available (41). As well as incorporating the effective energy differences, we need to implement within the simulation a representative pattern of cell types. In the *Drosophila* heart, Tin-positive cardioblasts express Fas3 at high levels and Svp-positive cardioblasts express Ten-m. We translate this into different adhesion energies per unit length between Tin-positive cardioblasts (ϵ_*TT*_), Svp-positive cardioblasts (ϵ_*SS*_), and between Tin- and Svp-positive cardioblasts (ϵ_*ST*_), similar to the outline given in Fig. 1B-E and Fig. 2D.

#### Initial configuration of the cells and initial cell alignment

By the end of dorsal closure, cardioblasts are brought into close contact. To initiate the cellular arrangement, we assume that the cells at the two ends of the heart are perfectly aligned and are at their resting length initially. To model geometric disorder, we take the length of the apical surfaces for Tin-positive and Svp-positive cardioblasts as *L* _0,*T*_ and *L* _0,*S*_ respectively. These lengths are simulated as Gaussian variables of mean 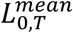 and 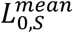 with standard deviation 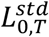 and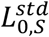, extracted from experimental quantification of cell size (Fig. 2F). Importantly, the experimentally measured rest lengths are used in the simulations, hence not all cells reach the final size. Given the tissue is constrained *in vivo* (51), these rest lengths represent the balance between the cell intrinsic rest length and external compression. We construct the simulated heart as two rows of cells formed by a succession of four Tin-positive cells and two Svp-positive cells patterns, with a total number of N_1_ =52 cells (number of cells in a heart row, Fig. 1B and 2G). Each cell length is picked as a random variable according to the rest lengths distributions (Fig. 2F). However, we constrain the total lengths of the two rows to be identical (Methods).

#### Initial cell matching without adhesion energy

We first consider the question: are the boundary constraints alone sufficient to ensure robust cell matching? To answer this, we take initial conditions that mirror experimentally measured cardioblast size and position (Fig. 2F-G) but with no adhesion energy (*i* .*e*. ϵ_*TT*_ = ϵ_*SS*_ = ϵ_*ST*_ = 0), Fig. 2D. As we take cells at their initial rest length, no equilibration is needed, and we calculate the mismatch of the initial condition. We performed 30 simulations per condition with confinement of different sizes and calculated the ensemble mismatch in cell alignment. This resulted in a mismatch increasing with system size, Fig. 2H. Interestingly, we find that at 52 cell length (the experimental size of the heart) we get a mismatch around 0.3, which corresponds to the mismatch value experimentally observed in the *Svp* ^*-/-*^ mutants where all cells are of the same type, Fig. 2C. In summary, geometric disorder, induced by cell size variability, is too large to enable precise cell matching merely by boundary constraints.

#### Calculating the mismatch with differential cell adhesion

We next investigated how the strength of selective adhesion impacts alignment. We first explored matching variations with only one cell type adhering, *i* .*e*. we varied ∈_*TT*_ alone with ∈_UU_ = 0 ∈_*ST*_ = 0, Fig. 3A-C. We computed mismatch evolution as a function of simulation iterations and took its value once it reached a steady-state value, Fig. 3C. The mismatch for each simulation was then averaged over 30 simulations to get the average final mismatch per condition. We find that the final mismatch decreases with increasing adhesion level.

**Fig. 3:**
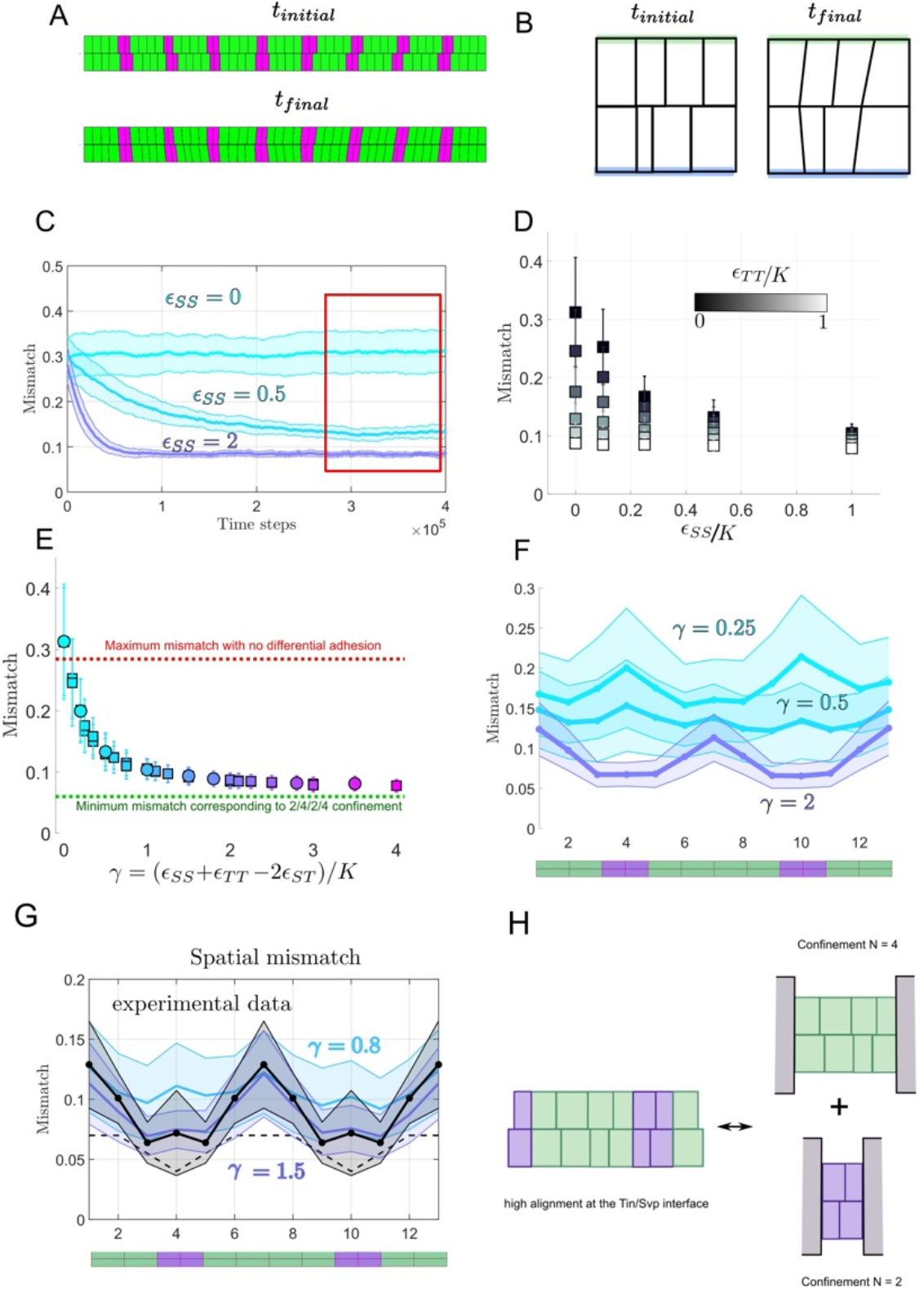
The interplay between differential adhesion, cell compressibility and cell size variability determines cell-matching. (A) *t* _*initial*_ corresponds to the initial configuration randomly sampled for 52 cells (4 Tin/2 Svp alternated pattern) and *t* _*final*_ corresponds to the cells configuration after convergence of mismatch. (B) Cell lengths have changed on the cell/cell interface while we still represent their initial length and position on the side not touching the interface (blue and green interfaces). (C) Ensemble averaged (N_2_ = 30 simulated embryos) mismatch evolution for different values of ε_SS_ and K = 1. All mismatches are calculated at steady state. (D) Final mismatch for different values of ε_SS_ and ε_TT_. (E) Final mismatch for different values of γ. Circles correspond to the case ε_ST_ = 0 and squares to ε_ST_ ≠0. (F) Spatial variation of mismatch across different cell types as depicted in the below cartoon (green = Tin positive cardioblasts, magenta = Svp positive cardioblasts.). (G) Spatial variations of the mismatch in the wildtype (in black) and comparison with the simulation (blue, purple). (H) Sketch showing how differential adhesion aligns heterogeneous cell-cell contacts, which generates local confinement of cells.

We then explored two cell types with selective adhesion by using different combinations of ∈_*SS*_ and ∈_*TT*_ with ∈_*ST*_ = 0, Fig. 3D. In general, increased adhesion energy led to reduced mismatch. Different combinations of ∈_*SS*_ and ∈_*TT*_ can lead to the same alignment. By systematically varying ∈_*SS*_, ∈_*TT*_ and ∈_*ST*_, we see that the equilibrium mismatch is determined by the parameter

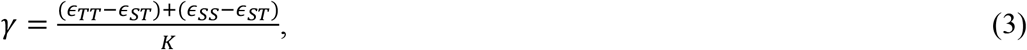

representing the competition between cells due to differential adhesion and cell compressibility, Fig. 3E. Here, γ corresponds to the typical length change that adhesion differences can generate for a single cell. For γ = 0μ*m*, there is no energy cost for different cell types to have a contact interface. The mismatch corresponds to the random case simulated in Fig. 2H; similar to the *Svp* ^*-/-*^ mutant, where all cells are of the same type. Mismatch decreases with increasing γ, reaching a plateau around mismatch of ∼ 0.07, dashed line in Fig. 3E. This implies that, beyond a certain level, increasing differential adhesion only weakly improves matching.

We measured spatial variations of the mismatch for different values of γ, Fig. 3F. We find that the mismatch is minimal at the Tin-Svp interface, as any contact between different cell types has some adhesion energy cost for γ > 0μ*m*, Fig. 3G. Alignment of cells inside a block of homogeneous cell types in the high γ limit is determined by the boundaries imposed by the highly aligned cells at the interface between different cell types. Within a region of Tin-positive cells, the level of cell mismatch converges towards the non-adhesive random case with N = 4, Fig. 3H. The mismatch between Svp-positive cells is similar to the random adhesion case with N = 2. The geometric variability in the lengths of the cell leading edges means that cells never reach zero mismatch even at very high adhesion values. Interestingly, for small values of γ, the pattern of cell mismatch is inverted, as geometric variability is higher for Svp cells.

Comparing the spatial profile of cell matching to the experimental data, we see that for γ > 0.5μ*m* the mismatch profile agrees well with experiment, Fig. 3G. Therefore, simple energy considerations, with a single fitting parameter γ, are sufficient to reproduce the wildtype matching phenotype.

### Energy scales are just sufficient to ensure robust matching

We do not know the specific effective energy levels associated with different adhesion interactions. To relate the values of ϵ_*TT*_, ϵ_*SS*_ and ϵ_*ST*_ to the underlying dynamics, we assume that they linearly depend on the energy scales of homophilic interactions of Fas3 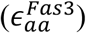 and Ten-m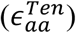, *e* .*g* .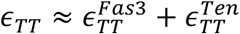. In our model, if there is no differential expression pattern of Fas3 and Ten-m then γ = 0 μm; *i* .*e*. there is no effective adhesion energy differences between cells. For *svp* ^-/-^ mutants (where all cardioblasts are Tin-positive) or the double mutant *Fas3* ^*-/-*^ ; *Ten-m* ^*-/-*^ (Fig. 4A) we expect γ ≈ 0.

**Fig. 4:**
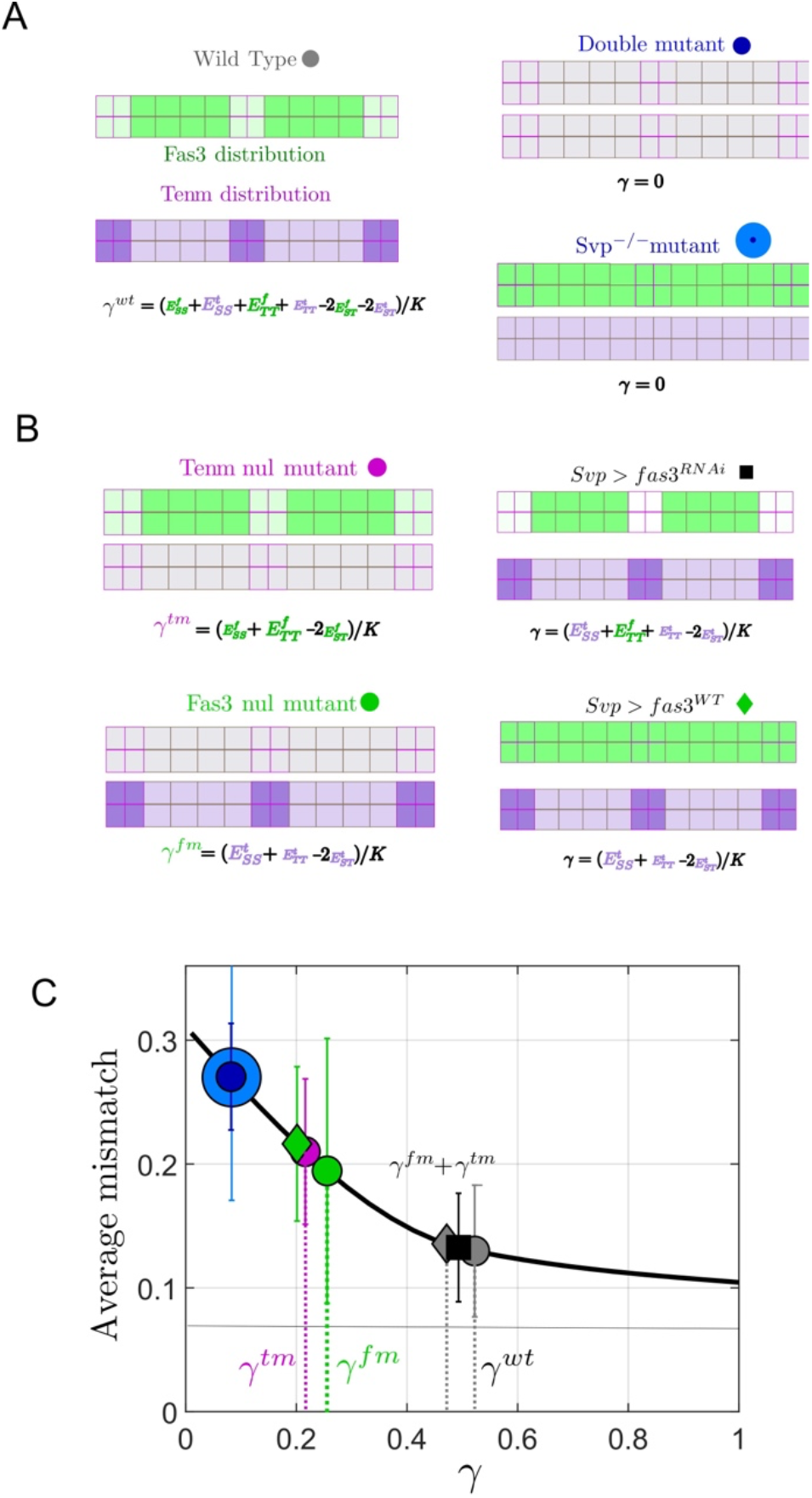
Equilibrium energy description replicates experimental observations in wild type and mutants. (A) Distribution of Fas3 (green) and Ten-m (purple) in wildtype embryos and the associated formula for γ. (B) Cartoons of relative interactions in different genotypes: wildtype (grey circle); *fas3* ^*-/-*^ *ten-m* ^*-/-*^ (blue small circle); *svp* ^*-/-*^ (blue circle); *fas3* ^*-/-*^ (green circle); *ten-m* ^*-/-*^ (purple circle); Svp-Gal4, Fas3-UAS (green diamond); and Svp-Gal4 > Fas3-RNAi-UAS (black square). Each adhesion energy can be decomposed into two contributions from Tin and Svp positive cells, leading to an approximation of γ. (C) Distribution of Fas3 and Ten-m mismatch in different genotypes (as defined in (B)) overlayed on the master mismatch curve of Fig. 2E. By placing the mismatch values on the curve, we can infer the corresponding effective γ on the *x* axis, with error bars representing the estimated error.

From previous work, the mismatch in a range of mutant conditions – such as disruption of Fas3 and Ten-m - is quantified (41). We utilized the results from Fig. 3E to estimate γ for wild-type and mutant conditions (Fig. 4B). We subsequently plotted the mismatch based on the inferred value of γ in a range of conditions (Fig. 4C). Can our model explain the observed values for γ in the mutant conditions?

From Fig. 4C, we have for the wildtype, γ_*WT*_ ∼ 0.5 μ*m*, which is smaller than the value γ_*WT*_ ∼ 1.0 μ*m* found by fitting the experimentally measured spatial variation of mismatch (Fig. 3G). However, this difference could be explained by the shallowness of the mismatch curve for γ > 0.5 μ*m*. Assuming that there is no interaction between Fas3 and Ten-m, we expect that γ_*WT*_ = γ_*Fas* 3_ +γ_*Tenm*_. We use the measured matching in *Fas3* ^*-/-*^ and *Tenm* ^*-/-*^ mutants to estimate the contributions to γ_*Tenm*_ and γ_*fas* 3_ respectively. From this, we find that γ_*Fas* 3_ +γ_*tenm*_ ∼γ_*WT*_ ∼0.5 (Fig. 4C).

To further test these assumptions, we considered additional mutant conditions. Overexpression of Fas3 in Svp cells should give similar value of γ to the *Fas3* ^*-/-*^ mutant, as in both cases the differential expression pattern of Fas3 is largely lost (Fig. 4B). As shown in Fig. 4C this is indeed the case. If Fas3 is reduced in Svp-positive cells, we expect ϵ_*SS*_ is reduced compared to wildtype. However, ϵ_*ST*_ is also likely decreased as direct interactions between filopodia with Fas3 is reduced between different cell types. Therefore, we do not expect γ to differ significantly from wildtype. Consistent with this, we find a γ similar to the wild-type case showing that ϵ_*ST*_ is small compared to ϵ_*SS*_ and ϵ_*TT*_. We can map a range of observations in mutant embryos onto a single curved for the mismatch described byγ, suggesting our adhesion hypothesis is a reasonable approximation of the underlying dynamics driving cell matching.

### Robustness of cell matching to perturbations

Here, we probe the robustness of the system to perturbations in cell number by considering variations in cell type specification. Within wildtype embryo populations, we observe variability in the spatial pattern of cardioblast specification. For example, we have observed embryos with three and five Tin-positive cardioblasts in a sector, as well as a solitary Svp-positive cardioblast, and an additional one (Fig. 5). Even when cell number is perturbed, we observe that the boundaries between different cell types are typically well defined. To test the model, we implemented different initial patterns of cardioblasts but otherwise kept parameters identical to the wildtype scenario with γ = 0.5. The system equilibrates to a state with clear boundaries between the different cell types and cells deform to align boundaries. This demonstrates that having two complementary adhesion processes is robust to perturbations in the structure of the heart as it develops.

**Fig. 5:**
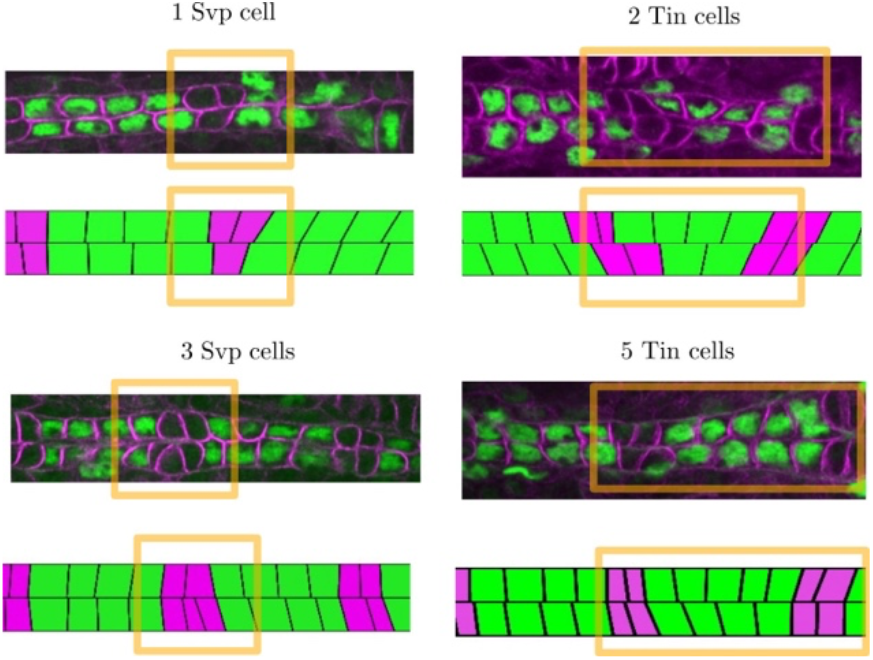
Matching correction of local defects. Various types of experimentally observed cell number defects in wild type embryonic hearts. Differential adhesion can partially compensate these defects by deforming cells in the range of γ ≈ 0.5 found in Fig.4.

**Fig. 6:**
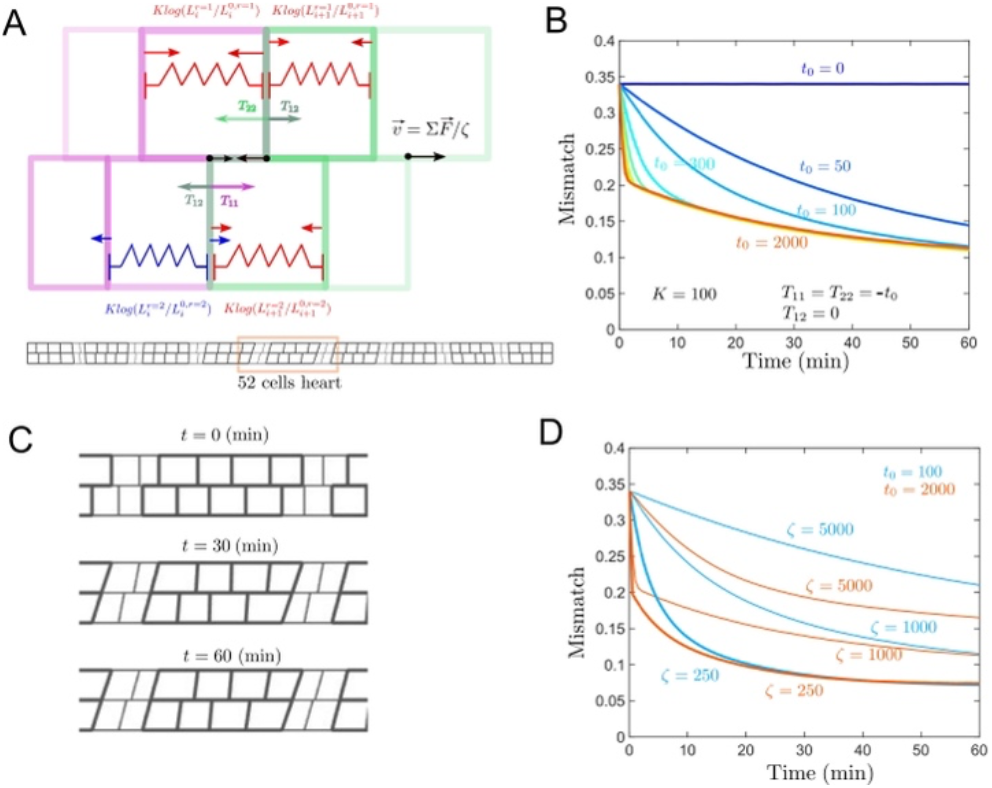
Simulating cell matching dynamics. (A) Vertex model implementation of interface dynamics. Each cell-cell lateral boundary on a row is defined by a vertex point. Elastic forces are represented in red/blue depending on if cells are stretched/compressed. Each cell-cell interface section between the two facing rows has a negative tension which acts on the vertices applying forces represented in magenta/green/grey depending on the interface type. The sum of these contributions divided by a friction coefficient, ξ, gives the vertex velocity. (B) We evolve the system (Methods) and compute the ensemble averaged mismatch over time (N = 30). We vary the tension *t* _0_ between 0 and 2000 while K is fixed to 100 and ξ = 1000. (C) Spatial alignment for 0 min, 30 min, 60 min for *t* _0_ = 2000, K= 100 and ξ = 1000 (D) Adhesion-independent cell alignment between cells of the same type depends on the friction coefficient ξ. Results shown for two different tensions, t_0_.

### Dynamic vertex model of cell matching

To evaluate how differential adhesion modulates cell matching dynamics, we used a vertex approach to model the time evolution of vertices at the interface between the two facing rows of cells, Fig. 6A (see Methods). By analogy with the equilibrium energy model used previously, vertices are submitted to elastic forces corresponding to effective elasticity and adhesion forces in the cell. We assume that adhesion produces a negative tension, which extends cell-cell contact interfaces. The sum of these contributions results in a force which, when divided by an effective friction coefficient, gives the vertex velocity. We explore the simple case of equal tensions, *T* _*SS*_ = *T* _*TT*_ = −*t* _0_ and *T* _*ST*_ = 0 with a fixed *K* and friction coefficient, ξ. Varying *t* _0_, keeping other parameters fixed, shows a lower limit to cell matching accuracy, as the time evolution of cell matching collapses onto a single curve for large *t* _5_, Fig. 6B. This limit is consistent with our equilibrium model. At high adhesion differences, the cell mismatch first undergoes a rapid decrease, which is adhesion-dependent. This corresponds to the phase of heterogeneous cell types boundaries shrinking. Below a mismatch of ∼0.2, the boundaries between cell types are perfectly aligned, Fig. 6C. The mismatch between cells of the same type decreases with slower, adhesion independent, dynamics that are friction dependent, Fig. 6D. Overall, we see that dynamically incorporating differential adhesion gives similar results to our equilibrium approach, supporting our above assumptions.

## Discussion

Even though biological systems are inherently out-of-equilibrium, we have demonstrated that an equilibrium energy argument is consistent with experimental observation of cell matching in the heart. This is consistent with the rapid dynamics of filopodia compared to the time of heart cells final alignment. The filopodia dynamics relate to the system effective temperature. The idea of effective temperatures in developing systems has also recently been used to describe jamming transitions during zebrafish development (52, 53). Our model shows that cell matching results from the interplay between cell-cell interactions (mediated by filopodia and specific adhesion molecules) and cell deformability. According to our model, the level of differential adhesion found in the wildtype heart is close to the minimal level required to obtain a well-aligned heart as the value γ = 0.5μm corresponds to the point where the mismatch curve in function of γ starts to flatten. We find that Fas3 and Ten-m contribute almost equally to the matching process. Furthermore, we demonstrate than increasing differential adhesion indefinitely does not accelerate the matching process beyond a certain point as the final alignment phase is adhesion independent.

A key result of our model is that short-ranged constraints (*i* .*e*. adhesion differences between two cell types) can propagate long-ranged order. We show that generating periodic segments of different adhesion molecules (i) aligns these segments at their interfaces and (ii) alignment inside each segment is due to geometric confinement and decreases with the segment size. Most previous theoretical work on the role of differential adhesion in tissue shaping has focused on how two (or more) distinct tissues can define clear boundaries (25). Our results are complementary to such work, and demonstrate that differential adhesion is a viable mechanism for aligning cells within a single tissue. Recent work has also highlighted how morphogen patterning of tissues coupled with mechanical feedback can build long-ranged tissue architecture (54). It will be interesting to explore how spatial information and mechanical feedback interplay in different tissues to ensure tight domain formation.

Here, we have not considered long-ranged actin structures that span multiple cells. These have been observed during heart closure, and appear to play an important role in cardioblast migration (46). Such structures could also play an important role in integrating mechanical information during closure.

Our work has potential application to other biological systems. During neurogenesis - where precise connections between neurons are required - Notch signaling (involve in lateral inhibition) has recently been shown to regulate neuronal wiring(13). Formation of the vasculature requires cell migration to precise locations and this is mediated by filopodia protrusions and lateral inhibition (55). During wound healing, cells need to repair wounds by forming precise connections. However, how processes at a single cell level are integrated to ensure long-ranged precise cell matching remains an open question (56). In our model, we do not consider heterogeneities in the mechanical properties. In other systems such as the brain, tuning of tissue stiffness ensures precise cell matching (57). More generally, coordinated cell migration can be directed by tissue stiffening (58). It would be possible to test the role of mechanical feedback in our model by adding viscoelastic relaxation of the cell lengths within a vertex model framework as described in Fig. 6.

## Author Contributions

S.T. and T.E.S. designed the project with input from S.Z. S.T. and M.S. performed the simulations and modelling. S.Z. provided all experimental data. S.T. and T.E.S. wrote the manuscript, with contributions from M.S. and S.Z..

## Acknowledgements

This work was supported by the University of Warwick, EMBO Global Investigator, Singapore Ministry of Education Academic Research Fund Tier 2 grant (MOE2018-T2-2-135), Singapore National Research Foundation Fellowship (NRF2012NRF-NRFF001-094) and HFSP Young Investigator Grant (RGY0083/2016), all awarded to TES. Tinman antibody was a gift from Manfred Frasch.

